# Implementation of an enriched membrane protein carrier channel for enhanced detection of membrane proteins in mass spectrometry-based thermal stability assays

**DOI:** 10.1101/2023.12.17.571949

**Authors:** Clifford G. Phaneuf, Alexander R. Ivanov

## Abstract

In this work, we developed a membrane-enriched stable isotope isobaric-labeled carrier channel (meSIILCC) for mass spectrometry-based thermal stability assay (MS-TSA). A proof-of-concept study demonstrated that the meSIILCC method could modestly improve membrane protein (MP) detection in MS-TSA experiments. An enhancement of 10% in identifications of membrane-proteins was observed in the meSIILCC group. Hydrophobicity analysis of the identified and quantified peptides using the grand average of hydropathy index confirmed the meSIILCC approach enriched for peptides of higher hydrophobicity characteristic of membrane-associated proteins.

To further improve meSIILCC, four membrane-protein enrichment approaches were compared. Using the selected and optimized workflow that utilized isobaric labeling-mass spectrometry, 8,662 protein groups were quantitatively characterized and then annotated based on their subcellular localization. The corresponding reporter ion intensities were used to construct a heatmap, which revealed an increased representation of proteins corresponding to the “plasma membrane” gene ontology term.

In a separate DMSO-only MS-TSA experiment, the optimally performing meSIILCC was added at 10-fold the protein content of the lowest heated aliquot from the MS-TSA, and isotope interference was found to be the highest in the 134N channel, while to a much lesser degree in other channels that were left empty.

To further assess the performance of meSIILCC in the DMSO-only MS-TSA experiment, an over-representation analysis was performed, which demonstrated that proteins exclusive to the meSIILCC group had more than a five-fold increase in gene ontology cellular component terms related to the “membrane” term.

We found 496 proteins from the DMSO-only MS-TSA experiment, which were identified across all replicates and shared between the meSIILCC and control that were annotated with “plasma membrane.” A close to 28% increase in the set corresponding to unique peptides was realized, using the meSIILCC approach, with a median value of 6.3 peptides per protein, compared to 4.7 in the control.

## 3.2 Introduction

Membrane proteins (MPs) are essential mediators of interaction between cells and their external environments. They have diverse roles in biology, such as critical regulators of cellular homeostasis^1^, but they can also be used as gateways for pathogens, as demonstrated in the studies of the SARS-CoV2 virus.^2^ The detection of MPs has essential implications in mechanism of toxicity, drug mode of action, and drug target identification studies.^3^ Only about 25% of the human proteome has been annotated as MPs,^4,5^ but these include the targets of >50% of all approved drugs.^6^ G protein-coupled receptors (GPCR) comprise the single largest class of approved drug targets (ADTs) and are predominantly located on the plasma membrane (PM).

Other large classes of MP identified as ADTs are nuclear receptors, ligand-gated ion channels, and ion-gated ion channels. MPs have historically been underrepresented in research because of their unique biophysical characteristics.^7–9^ Low abundance and poor aqueous solubility are common challenges associated with MP analysis. Despite these difficulties, MPs remain an attractive class of drug-targets pursued by the pharmaceutical industry.^10,11^

Thermal proteome profiling (TPP)^12–14^ was adapted from the cellular thermal shift assay (CETSA),^15^ and is now routinely used to study drug-protein interactions^16–18^. By leveraging the phenomenon of ligand-induced thermal stabilization of proteins, MS-TSA such as TPP and CETSA MS^19^ have enabled comprehensive target profiling in biologically relevant matrices for the identification of unknown targets and toxic mechanisms of action^3^.

Increased MP coverage in MS-TSA was successfully achieved after incorporation of a mild detergent,^20^ and further improvements, specifically targeting PM proteins, was demonstrated with the establishment of cell surface (cs)-TPP^21^. However, cs-TPP has yet to become widely adopted, presumably due to the complicated workflow, which combined TPP with selective enrichment of glycosylated cell surface proteins using a biotin-streptavidin pull-down system.

The use of a stable isotope isobaric-labeled carrier channel (SIILCC) technique has become a widely adopted approach to improve the detection of low abundant analytes, particularly in the area of single-cell proteomics.^22–25^ The SIILCC concept exploits the combined ion current effect associated with isobarically labeled peptides.^26^ Typically, one channel of a multiplex is designated as a “trigger channel” or “carrier channel” and comprised of the sample matrix at a several times higher total protein amount than the amounts in other channels. In a recent publication, we demonstrated an increase in drug-treated vs. control protein-level melting-curve comparisons using an SIILCC addition at four times the total protein concentration of the lowest heated sample.^27^

Here, we demonstrate the use of an <u>M</u>P-<u>e</u>nriched <u>SIILCC</u>, which we refer to as meSIILCC, to improve the quantitative detection of MP melting-curve comparisons. As a proof-of-concept study, we prepared a meSIILCC using a commercially available detergent-based extraction kit, and demonstrated increased quantitative identifications of MPs, increased overall quantitative identifications, and increased “Grand Average of Hydropathy” (GRAVY) scores. Next, several protocols were assessed for isolation of an MP fraction to be used as meSIILCC. A tandem mass tag (TMT) labeling experiment was used to compare multiple MP isolation techniques, which identified a commercially available cellular fractionation kit as outperforming other techniques and kits.

We further tested the use of a meSIILCC in a TMTpro 18-plex format by evaluating a ten-fold total MP level compared to the total protein concentration of the lowest heated sample (37 °C), and found isotopic impurities present in adjacent channels.

## Experimental

### Materials and Reagents

Halt™ Protease and Phosphatase Inhibitor Cocktail (100X), triethylammonium bicarbonate (TEAB), tris(2-carboxyethyl)phosphine (TCEP), tris(hydroxymethyl)aminomethane (Tris), sodium dodecyl sulfate (SDS), Titer Plate Shaker (model #4625), Veriti thermocycler, NP-40 Surfact-AmpsTM detergent, MicroAmp^TM^ optical adhesive film, FBS, formic acid (FA), Penicillin-Streptomycin (10,000 UmL-1), T225 flasks, 1x PBS, and all solvents at Optima LC-MS grade were purchased from Thermo Fisher Scientific (Waltham, MA) Jurkat E6.1 leukemic T-cell lymphoblasts were purchased from the American Type Culture Collection (ATCC, Manassas, VA). 2-chloroacetamide (CAA), RPMI-1640, and dimethyl sulfoxide (DMSO) were purchased from Sigma-Aldrich (St. Louis, MO). 96-well semi-skirted Hard shell PCR plates were purchased from Bio-Rad (Hercules, California). MultiScreenHTS GV Filter Plates, 0.22 µm (MSGVN2250), Benzonase nuclease, and CXCR3 inhibitor (SML0911) were purchased from MilliporeSigma (Burlington, MA). Thermomixer® C was purchased from Eppendorf AG (Hamburg, Germany). Trypsin/Lys-C mix was purchased from Promega (Madison, WI), Cytiva (Emeryville, CA, USA) Sera-Mag SpeedBeads™ Carboxyl Magnetic Beads (hydrophobic and hydrophilic) were purchased from Fisher Scientific. MEK inhibitor RO4987655 (MedChemExpress, 1 Deer Park Dr, Suite Q, Monmouth Junction, NJ 08852, USA)

### Cell Culture

Jurkat cells were expanded in media containing RPMI-1640 supplemented with 10% FBS and 100 UmL-1 Penicillin-Streptomycin in a 37 °C incubator with 5% CO2. Cells were cultured and maintained at a density of 1e^6^ cells per mL, and fresh media was exchanged every two days and the day before any experiment.

### Intact cell MS-TSA sample preparation

All samples were prepared in triplicate. Prior to experiments, cells were washed with 1x PBS twice and resuspended at a density of 50e^6^ cells/mL in 2 mL low-bind tubes. Either DMSO or a commercially available compound MEK inhibitor RO4987655, MedChemExpress, 1 Deer Park Dr, Suite Q, Monmouth Junction, NJ 08852, USA) was added where appropriate, which resulted in a final concentration of 10 µM compound and 1% DMSO. After a 1-hour incubation at 37 °C, 80 µL aliquots of each suspension were transferred to 96 well platesand heat-challenged at eight temperature points (37, 41.5, 46, 50.5, 55, 59.5, 64, and 68.5 °C) for 3 minutes in a thermocycler and allowed to equilibrate to room temperature on the benchtop for 3 minutes. To each well, NP-40 detergent was added to a final concentration of 0.4% and mixed by pipetting. Samples were snap-frozen in liquid nitrogen for 2 minutes and thawed in a room temperature water bath at room temperature for 2 minutes for a total of four cycles and stored at −80°C. A mixture of nuclease, protease, and phosphatase inhibitors was added to each sample for a final dilution concentration of 25 UmL-1 and 1x, respectively, and incubated at 4 °C for 20 minutes on a titer plate shaker at half maximum speed. The insoluble fraction of the lysate was cleared by filtration using 0.22 µm filter plates. Subsequently, 80 µL of supernatant from each well was transferred to a new set of 8-well PCR strips and stored on ice. Total protein concentration was estimated using BCA assay according to the manufacturer’s protocol, and samples were stored at −80°C before further processing.

### MP pellet preparation using Dounce extraction

An MP pellet was prepared according to previous work^28^ with minor modifications. For each MP pellet, 100e^6^ Jurkat cells were harvested and washed twice with ice-cold PBS. The cells were pelleted at 400 x g for 10 minutes at 4°C and then resuspended in 5 pellet volumes of hypotonic buffer (10mM HEPES pH 7.9, 10mM KCl, 1.5mM MgCl2, 0.5 mM DTT, and supplemented with HALT protease and phosphatase inhibitors). The suspension was incubated on ice for 10 minutes, pelleted again, and resuspended in two pellet volumes of hypotonic buffer. The suspension was then transferred to a 2 mL Dounce homogenizer (Bellco Glass Inc. Vineland, New Jersey, USA), and cells were lysed using 30 strokes with the “type B” tight-fitting pestle on ice. The lysate was transferred to a 1.5 mL low bind tube and centrifuged at 1000 x g for 10 minutes at 4 °C to remove unbroken cells and debris. The supernatant containing membranes was then transferred to thick-walled polycarbonate centrifuge tubes and centrifuged at 100,000xg for 1 hour at 4 °C. The supernatant was removed, and the tube containing the pellet was washed with 0.5 mL of hypotonic buffer, followed by another 15-minute centrifugation at 100,000xg at 4°C. The rinsate was removed, and pellets were stored at −80°C until used.

### MP pellet preparation using Mem-per plus kit

Samples were prepared according to manufacturer’s protocol with slight modifications. Briefly, cells were washed with Cell Wash Solution (CWS) and centrifuged at 300 x g for 5 minutes. The supernatant was removed, the cells were resuspended in CWS, and 1.5 mL aliquots containing 10e^6^ cells were transferred to 2 mL low bind tubes. The samples were then centrifuged at 300 x g for 5 minutes, and the supernatant was discarded. Each pellet was then resuspended in 0.75 mL of Permeabilization Buffer, briefly vortexed, and incubated for 10 minutes at 4 °C with constant mixing. The samples were then centrifuged at 16,000 x g for 15 minutes. The supernatant containing the cytosolic proteins was removed, and 0.5 mL of solubilization buffer (containing protease and phosphatase inhibitors) was added to each tube, and the pellet was suspended by pipetting up and down. The samples were then incubated at 4°C with constant mixing for 30 minutes, followed by centrifuging at 16,000 x g for 15 minutes at 4°C. The supernatant containing the membrane protein fraction was then transferred to a new low-bind tube and frozen at −80°C until use.

### Control sample preparation

Samples were prepared similarly to those in the intact cell heat treatment, with the exception that the heating step was omitted.

### MP pellet preparation using the Invent Biotech membrane isolation kit

Samples were prepared according to the manufacturer’s protocol with slight modifications. The filter cartridges were placed in collection tubes and stored on ice. For each sample, 50 x 10e^6^ cells were washed twice with ice-cold 1xPBS and pelleted by 300 x g for 5 minutes. The supernatant was removed, and the pellet was resuspended in 500 μl buffer A, followed by a ten-minute incubation on ice. The tubes were then vigorously mixed for 30 seconds, and the suspension was immediately transferred to the filter cartridge and capped. The cells were lysed by passing through the filter cartridge in the centrifuge at 16,000 X g for 30 seconds. The filter was discarded, and the pellet was resuspended by vigorously vortexing for 10 seconds. Intact nuclei were removed from the suspension by centrifugation at 700 X g for one minute, and the supernatant was transferred to a new 1.5 mL tube. The cytosolic and total membrane fractions were separated by centrifugation at 16,000 x g at 4 °C for 30 min. Half of the resulting pellets, which contained the complete membrane (iCM) fraction (plasma and other intracellular membranes) were stored at −80°C until ready to use. The remaining pellets were resuspended in 200 μL of buffer B, and centrifuged at 7,800 X g for 5 min at 4 °C. The resulting pellet, which contained organellular membranes, was discarded, and the supernatant was transferred to a new 2.0 ml low-bind tube. To each sample 1.6 mL cold 1x PBS was added and mixed. The suspensions were then centrifuged at 16,000 X g for 30 minutes, and the supernatant was discarded. The resulting pellets, which contained purified plasma membranes (iPM), were stored at −80°C until ready to use.

### Digestion and Sample Processing

Pellets were thawed and reconstituted in 1% SDS. For heat-challenged samples, equivalent volumes from each heat-challenged temperature point that corresponded to 25 µg from the 37°C aliquot of each replicate were transferred to 96 deep-well plates. MP-enriched pellets to be used for SIILCC were also reconstituted in 1% SDS and processed similarly in aliquots of 12.5 µg.

Tris was added to each sample to a final concentration of 100 mM at pH 8, and each sample was brought to a volume of 106.25 µL with deionized water. Reduction and alkylation were performed in a one-step process using a final concentration of 20 mM TCEP and 40 mM CAA in each sample, followed by heating at 57 °C for 1 hour in a thermomixer. Detergent removal was accomplished using an SP3 magnetic bead approach^29^ using a Kingfisher automated purification system (Thermo Fisher Scientific). Samples were diluted 1:1 with 100% ethanol. Equal amounts of hydrophilic and hydrophobic magnetic beads were washed three times with DI water, and a 250 µgµL-1 slurry was prepared in 50% ethanol. Slurry aliquots of 100 µL were distributed to a 96-deep well plate and subsequently washed again for two minutes in 50% ethanol using the Kingfisher automated system. The magnetic beads were then transferred into the sample plate and incubated for 14 minutes with gentle mixing. The bead-protein mixture was washed three times with 70% ethanol and then transferred to a 350 µL 96-well plate containing 100 µL of 100 mM TEAB (pH 8.5). Trypsin/LysC mix was added to each sample at a ratio of protein to the enzyme of 50:1 (for heat-treated samples, this was based on the 37 °C sample) and incubated in the Thermomixer® C at 37 °C 500 rpm overnight. The next morning, a second aliquot of trypsin/LysC mix was added for a final ratio of protein to enzyme of 25:1, followed by a four-hour incubation. Beads were removed by filtration with a 0.22 µm filter plate, transferred to 8-well PCR strips, and stored at −80°C until used.

### Isobaric Labeling

Peptides were labeled with TMTpro (Thermo) according to the manufacturer’s recommendations with slight modifications. Briefly, digests were thawed and tested using pH strips to ensure the pH was between 8 and 8.5. The labeling reagents were allowed to equilibrate to room temperature and solubilized with anhydrous acetonitrile for 5 minutes with occasional vortexing. Each TMT reagent was added to the appropriate sample at a ratio of label to the total peptide of 8:1 and a final acetonitrile concentration of 19.5%. Incubation was carried out at room temperature for 1.5 hours in the dark and quenched with 0.3% hydroxylamine final concentration for 15 minutes. The labeled digests were pooled accordingly.

### Offline high pH fractionation

Desalted peptides from the sample set prepared for fractionation were subjected to offline high-pH reversed-phase separation on an Agilent 1290 HPLC using a Phenomenex XBridge BEH C18 Column, 130Å, 3.5 µm, 3 mm X 150 mm (Waters, MA), which was heated to 45 °C. The dried desalted isobarically-labeled digested samples were resuspended in 100 µL of solvent A (20 mM ammonium formate, pH 10, and 5% acetonitrile), and the entire sample was injected onto the column. The separation was accomplished using the following gradient: 3% solvent B (20 mM ammonium formate, 95% acetonitrile, and pH 10) isocratic flow (450 µL/min) for 9 minutes, 3-35% solvent B in 40 minutes, 35-90% solvent B in 4 minutes, 90% solvent B isocratic flow for 3 minutes, 90-3% solvent B in 2 minutes, and 3% solvent B isocratic flow for 20 minutes. The resulting fractions were collected every minute and then concatenated into 24 fractions, dried in a Speedvac, and stored at −80 °C until ready to use.

### Nano-Liquid Chromatography and Mass Spectrometry Conditions

Nano-flow Liquid Chromatography coupled to Mass Spectrometry (nLC-MS) analysis was conducted using either an Evosep One (Evosep, Odense, Denmark) for the POC MS-TSA pilot and optimal meSIILCC selection, while a Vanquish Neo UHPLC system (Thermo Scientific, Waltham, MA) was used for the DMSO-only MS-TSA experiment. In both cases, chromatographic systems were interfaced with an Orbitrap Exploris 480 mass spectrometer (Thermo Scientific, Waltham, MA).

The Evosep one was equipped with an EV1106 (15 cm x 150 µm) separation column using the Extended Method (88 min.). Chromatography with the Vanquish Neo was accomplished using a PepMap™ Neo Trap Cartridge (5 mm x 300 µm, 5 µm particle size) in line with an IonOpticks (Melbourne, Australia) Aurora Elite column (Gen 3, 15 cm x 75 µm). Samples were reconstituted in 0.1% FA, and the total peptide concentration was determined by the Pierce™ Quantitative Peptide Assays (Thermo Scientific, Waltham, MA). One µg of each sample was loaded on the trap column and washed for 5 minutes at maximum pressure (1,500 bar) with solvent A (0.1% FA in MS-grade water). The reversed-phase gradient started at 1% solvent B, which consisted of 0.1% FA in mass spectrometry-grade acetonitrile and held constant for 5 minutes. Solvent B was increased to 3% in one minute and to 18% over the next 60 minutes. Solvent B was further increased to 30% over the next 20 minutes A washout period occurred by increasing solvent B to 80% in 3 minutes and held constant for two minutes. The analytical column was re-equilibrated with ten column volumes of solvent A.

MS1 scans were acquired in positive mode (2,300 V) using a resolution setting of 60,000. Normalized automatic gain control (ACG) target was set at 300%, with a maximum injection time of 25 ms. Peptides with a charge state of 2-6 in the 400-1600 m/z range that had a minimum intensity threshold of 5,000 were selected for fragmentation using a 1.6 m/z widow. MS2 scans were acquired in centroid mode using a cycle time of 1 second, with a resolution setting of 30,000. The normalized collision energy setting was 35, normalized automatic gain control (ACG) target was set at 200%.

### Data analysis

Raw files for each replicate were first analyzed as fractions using Proteome Discoverer (PD, version 3.0 Thermo Fisher Scientific) with the Sequest HT search engine. Tryptic peptides were included in the searches with a maximum of two missed cleavages. Precursor and fragment mass tolerances were set to 15 ppm and 0.02 Da, respectively. Static modifications included TMTpro on the peptide N-terminus and lysine residues and carbamidomethylation of cysteine residues.

Oxidation of methionine residues was set as a dynamic modification, along with acetylation of the protein N-terminus. The UniProt human database, including reviewed reference proteome release 2018_03 or 2022_01 was used for the POC experiment or others, respectively. Peptide filtering was set to “high” confidence, resulting in a 1% FDR.

Melting curves and melting temperature(s) (T_m_) were determined using TIBCO Spotfire^TM^ (TIBCO, version 11.4), or an in-house R-based script using the previously described nls2 package^30^. An in-house virtual basic (Microsoft Corporation, 2023) script was first used to pre-process replicates. Each replicate treatment (“DMSO” or “TREATED”) was normalized to the 37 °C channel. Corrected ratios were imported into Spotfire (TIBCO, version 11.4), and the T_m_ for each replicate treatment was determined using the “logistic regression curve fit” function.

## Results and Discussion

### A proof-of-concept pilot indicates meSIILCC benefits MP identifications

An initial proof-of-concept (POC) study was conducted using the previously described^27^ dual specificity mitogen-activated protein kinase kinase 2 inhibitor (MEK2) model system to rapidly assess the use of meSIILCC for improved detection of MPs in MS-TSA experiments. The Mem-PER^™^ Plus Membrane Protein Extraction Kit (Thermo Scientific, Waltham, MA, USA) was used to prepare a meSIILCC that was subsequently digested and labeled with the 135N TMTpro reagent. The meSIILCC was pooled with other (Figure 1) differentially heated intact cell aliquots at four times the total protein amount of the lowest heated aliquot determined by the bicinchoninic acid assay (BCA assay, Thermo Scientific, Waltham, MA, USA).

**Figure 1.**
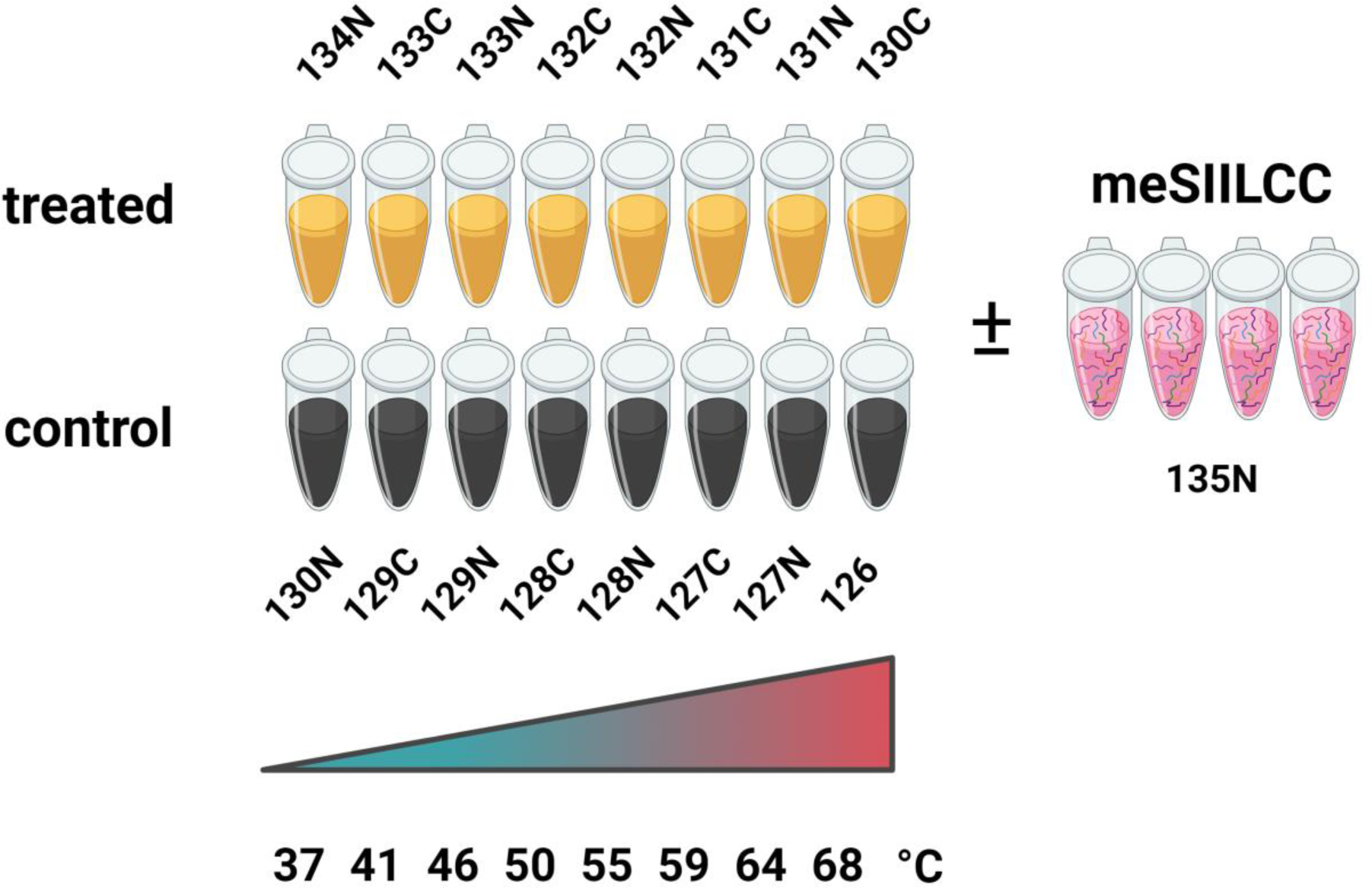
Isobaric labeling scheme for POC experiment using TMTpro 18-plex. The meSIILCC (135N) was included at 4-fold the total protein content of the lowest heated aliquot. Created with www.Biorender.com

We first compared protein groups, peptides, PSM identifications, and MS^2^ scans (collectively referred to as “proteomic identifications”), as shown in Figure 2. In the control group, 7,895 (±119) protein groups were identified, while 8,356 (±38) were identified in the meSIILCC group, resulting in a modest 5.8% improvement. The average count of peptide spectral matches (PSMs) for the control group was 169,655 (±10,426), compared to 188,396 (±7,133) in the meSIILCC group, demonstrating an 11.1% improvement. A 16. % improvement in peptide identifications was realized with 106,171 (±2,570) identified in the meSIILCC group in contrast to 91,022 (±5,633) in the control. As expected, the average number of MS2 scans for each group was similar; the control group averaged at 877,676 (±2,708), while the meSIILCC group averaged 868,686 (±19,172).

**Figure 2.**
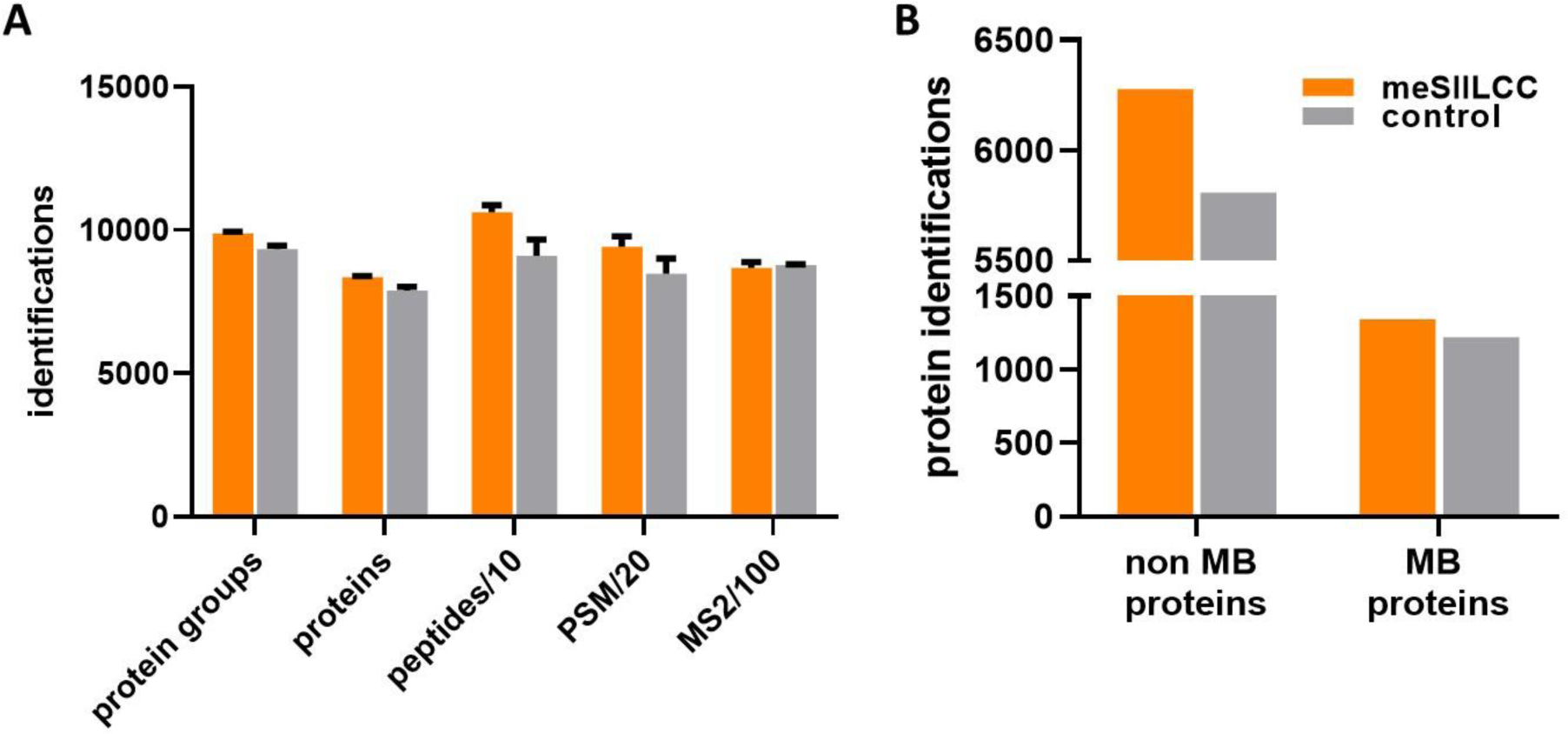
Identifications for the POC experiment compared to the control. **A)** Proteomic identifications. **B)** Protein identifications of MB and non-MB proteins.

We also considered the overlap and uniqueness of protein groups that were identified across all three replicates, as shown in the Venn diagrams of Figure 3. In the control group, 7,026 protein groups were consistently identified, and 7,619 in the meSIILCC, resulting in an 8.4% improvement of consistently identified protein groups when using the MB trigger. Additionally, 6,805 protein groups were commonly identified in both groups across all replicates.

**Figure 3.**
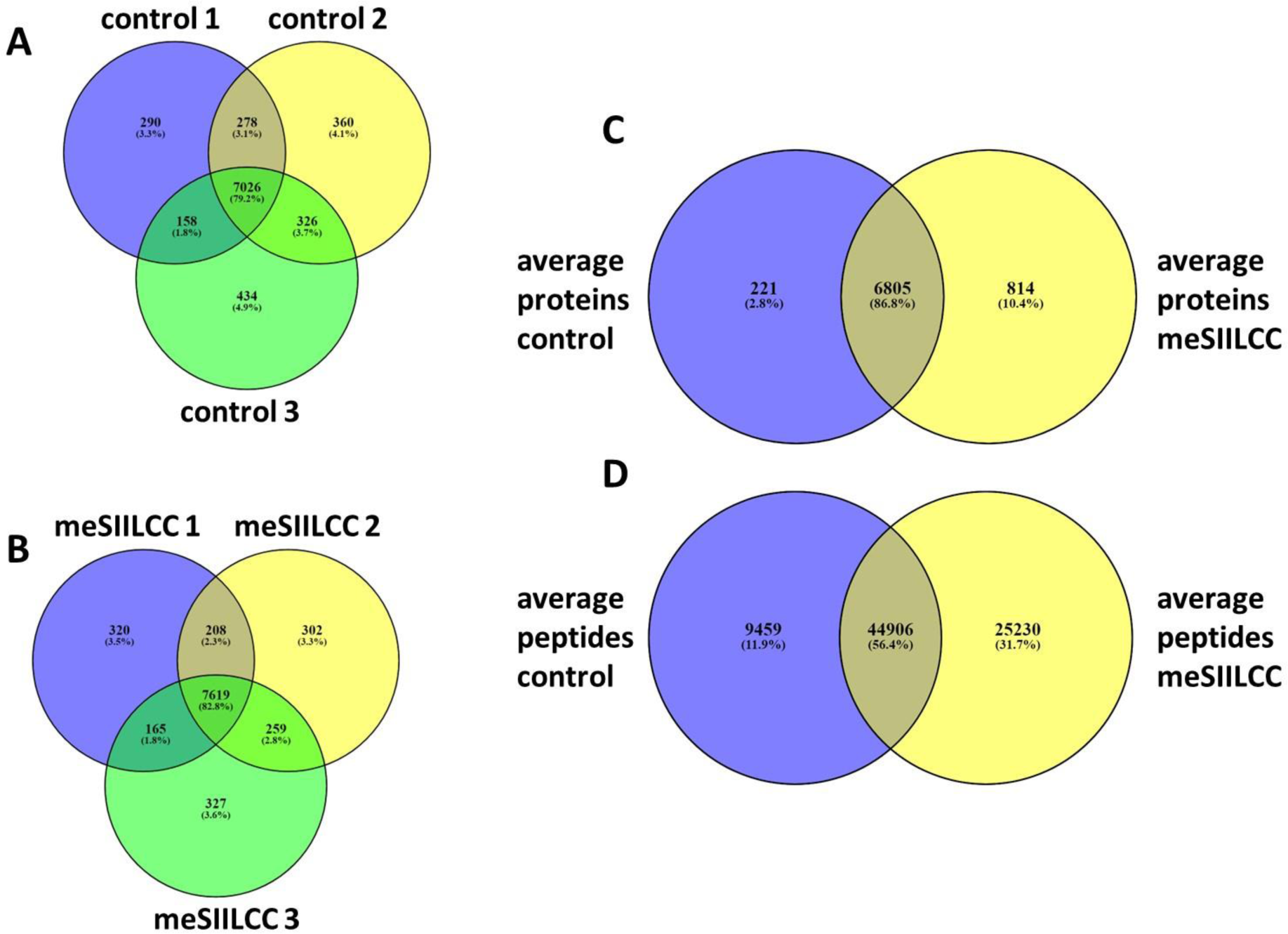
Overlap of identifications found in the POC experiment. **A)** Proteins identifications for control and **B)** meSIILCC replicates **C)** unique and common proteins, and **D)** peptides identified across all replicates.

**Figure 4.**
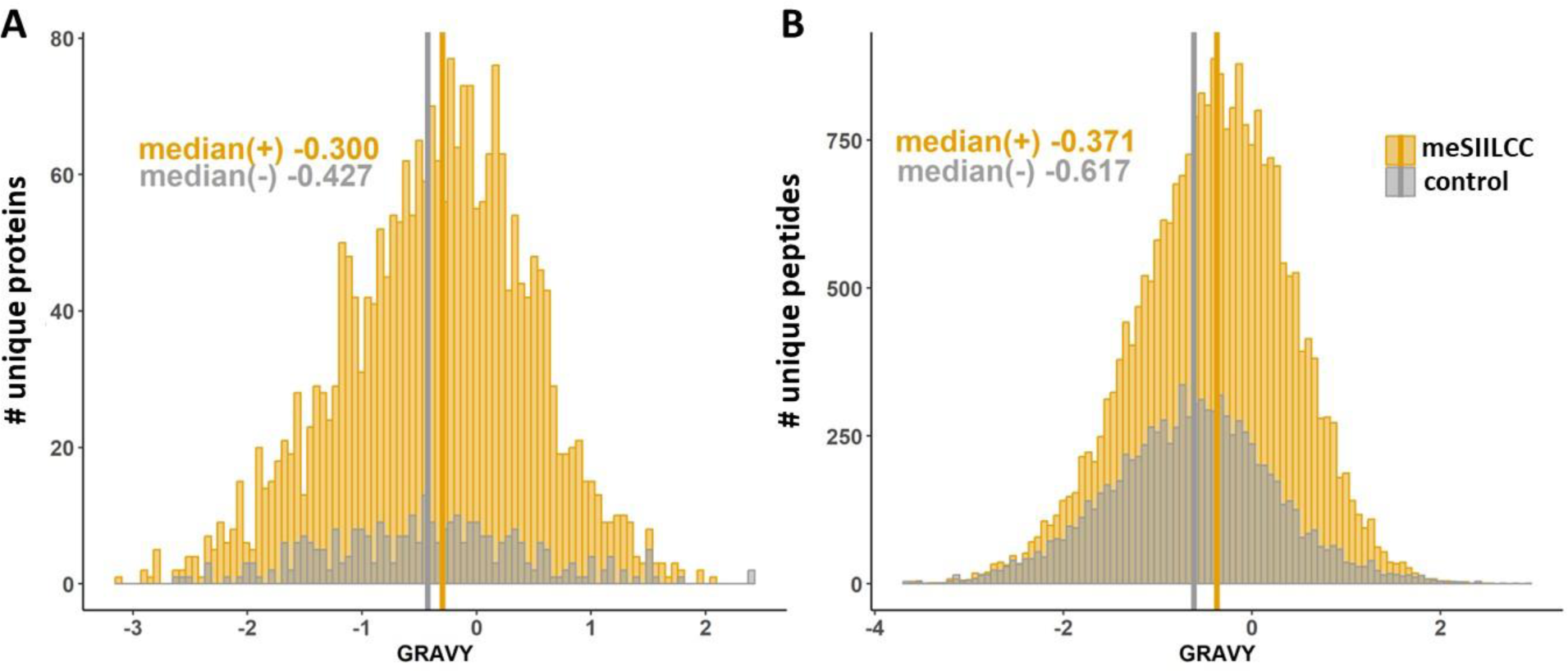
Distributions of GRAVY scores and their medians for control and meSIILCC in the POC experiment for **A)** Unique proteins and **B)** unique peptides which were detected across all replicates. Median GRAVY scores shown by bold vertical lines.

Using the Reviewed (Swiss-Prot) reference proteome from UniProt, which comprised 20,426 entries as of October 2023, we annotated proteins from both groups as either a MP or non-MP. If a protein contained either a transmembrane or intramembrane domain, suggesting its association with either plasma or organellular membranes, it was considered an MP. Proteins without either domain were classified as non-MPs. The control group contained 5,807 non-MPs and 1,219 MPs, while 6,278 non-MPs and 1,341 MPs were identified in the meSIILCC group (Figure 2B). Interestingly, this translated into an 8.1% improvement in identifications of non-MPs and a 10% improvement for MPs in the meSIILCC group compared to the control, indicating the meSIILCC prepared by the MEM-PER kit enriched for membrane and non-MPs.

We also explored the hydrophobic characteristics of peptides identified between the two groups, anticipating that the MP-enriched sample would yield peptides with greater hydrophobicity. The GRAVY index is a standard measure used to determine the hydropathy of a peptide or protein, based on the average score of its constituent amino acids. Utilizing the Kyte-Doolittle scale^31^, a positive GRAVY score is indicative of more hydrophobic molecules, while a negative score suggests more hydrophilic molecules. Only peptides consistently detected across three replicates in both experiments were included in the GRAVY analysis. Of these, 9,459 peptides were exclusive to the control group, yielding a median GRAVY of −0.617. In contrast, over 2.6-fold (25,230) were exclusively identified in the meSIILCC-exclusive group, with a median GRAVY of −0.371. The shift in GRAVY values suggests that peptides from the meSIILCC experiment were intrinsically less hydrophilic, a characteristic of membrane-associated proteins, which ensures their stability in the lipid-bilayer^32^.

Taken together, these findings encouraged us to further explore the use of the meSIILCC approach to improve MP detection in MS-TSA.

### A commercially available MP enrichment kit was selected to enhance MP extraction approach for preparation of meSIILCC

MPs are a heterogeneous group of molecules, having a wide range of hydrophobicities, distinct localizations within the lipid bilayer, and interactions with various cellular components^32,33^. These diverse attributes are underscored by the existence of numerous extraction protocols^7,34,35^. Although the importance of these proteins has been well-established, a unified isolation technique that addresses challenges such as contamination of co-localizing proteins, low extraction efficiency, poor reproducibility, and limited throughput has yet to be developed.

In order to further improve the performance of meSIILCC, four MP extracts were tested in total; the complete membrane (iCM) and plasma membrane fractions (iPM) from Invent Biotechnology’s Minute^™^ Plasma Membrane Protein Isolation and Cell Fractionation Kit (Invent Biotechnologies, Plymouth, MN, USA), the PM fraction prepared using the Mem-PER^™^ Plus Membrane Protein Extraction Kit (Thermo Scientific, Waltham, MA, USA) which is referred to as MEMPER, and the MP fraction prepared by Dounce homogenization ^28^ (Dounce).

We employed isobaric labeling-MS using TMTpro 16-plex to simultaneously quantify the four MP extraction profiles with a control, and quantitatively identified 8,662 protein groups. These values were used to construct a heatmap of log_2_ fold-change (log_2_FC), transformed reporter ion intensities as shown in Figure 5.

**Figure 5.**
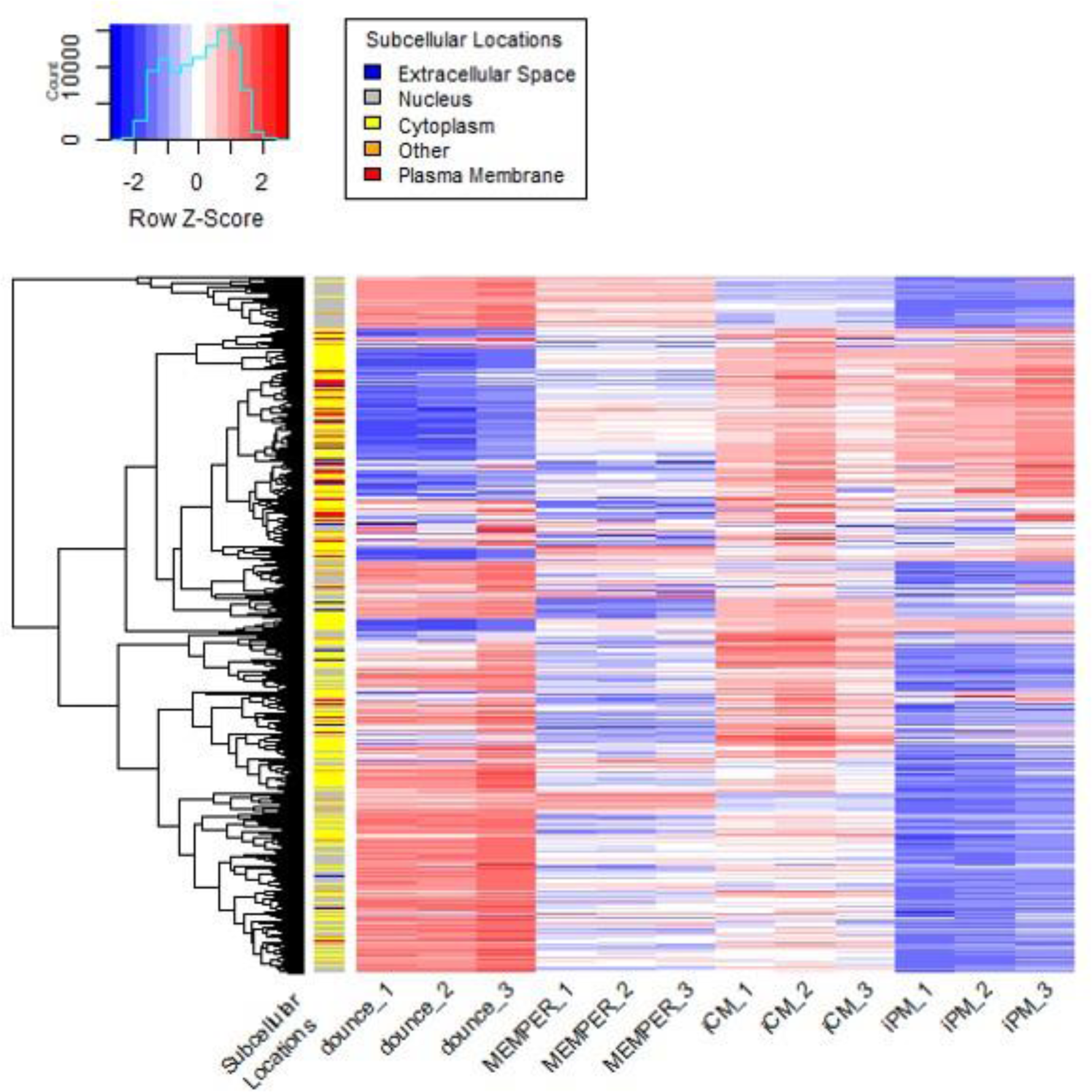
Heatmap annotated with subcellular locations which show log2-fold change transformed reporter ion intensities from triplicate MP extraction protocols.

The Swiss-Prot reference proteome described above was used in combination with protein subcellular locations from Qiagen’s Ingenuity Pathway Analysis (IPA, QIAGEN Inc., https://digitalinsights.qiagen.com/IPA).

IPA classifies proteins into five unambiguous categories: nucleus, cytosol, extracellular matrix, plasma membrane, and others.

We identified 3,996, 309, 2,645, 611, and 873 protein groups annotated as cytoplasm, extracellular space, nucleus, other, or plasma membrane, respectively. The number of protein groups annotated as “plasma membrane” for the mean Dounce homogenizer, MEMPER, iCM, and iPM was found to be 233, 406, 576, and 536, respectively.

The heatmap in Figure 5 illustrates a clustering of positive log_2_FC values along with PM-annotated proteins in the triplicate iPM extracts. Similar clustering was noted in the iCM extracts, in addition to the presence of positive log2FC features associated with subcellular locations other than PM, indicating that iCM extracts may not be a suitable choice for our planned MP-enriched MS-TSA experiments.

We found a distinct and almost opposite log_2_FC patterns in the heatmap between iPM and Dounce extracts, highlighting the unique selectivity of the respective enrichment procedures. Many log_2_FC features annotated with nucleus were shared between Dounce and MEMPER extracts, while Dounce extracts contained 70 unique proteins annotated as PMs (Figure 6A).

**Figure 6.**
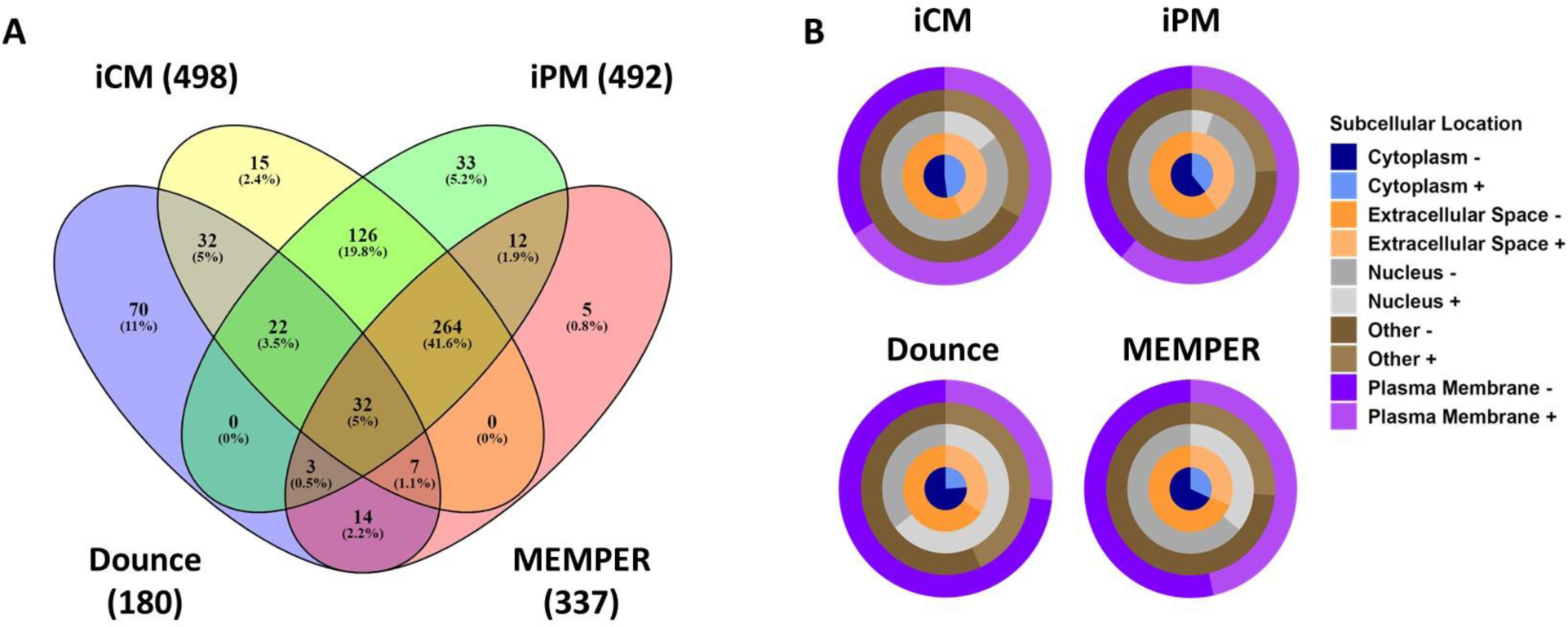
Comparison of PM-annotated protein identifications and log_2_FC distributions across subcellular locations in four MP-enriched extracts. **A)** Venn diagram showing the relationship of PM-annotated protein identifications. **B)** The distribution of positive and negative log_2_-fold changes for annotated subcellular locations.

To further discriminate the MP protein extraction effectiveness for the different evaluated protocols, the percentage of identifications for each subcellular location having a positive or negative log_2_FC was determined (Figure 6B). The Dounce protocol displayed the lowest percentage of positive log_2_FC values annotated with the “plasma membrane” cellular localization term (27%), while 47% of identifications in the MEMPER set had positive log_2_FC values. The iCM and iPM had 66, and 61% positive log_2_FC values annotated as “plasma membrane”, respectively.

The iPM extraction technique was selected for our subsequent experiments as a result of its combined performance to selectively enrich PM-annotated proteins, while at the same time excluding other proteins not annotated with PM. Notably, we observed positive log_2_FC values for many organelle MPs (OMPs) and speculate that our sample preparation technique using the selected commercial MP-enrichment kit requires further scrutiny if exclusively focusing on PM proteins in future studies.

### Reporter ion isotope interference was observed in empty TMT channels when using a meSIILCC at 10-fold the total protein of the lowest heated MS-TSA

Sigmoidal curve fitting of normalized reporter ion intensities is a common approach to determine melting temperatures (T_m_) for MS-TSA.^13,36,37^ However, isotope impurities within isobaric labeling reagents can be problematic when employing SIILCC.^38–41^ We assessed isotopic interference from meSIILCC by conducting triplicate MS-TSA experiments using only dimethyl sulfoxide (DMSO)-treated samples with and without meSIILCC. As shown in Figure 7, eight differentially heated intact cell aliquot digests were labeled with 126-130N TMTpro 18-plex reagents, while channels 130C-134C were not used. Pooled samples were split in two equal parts. A 10-fold excess of 135N-labelled meSIILCC (using the iPM enrichment) was added to one part, while the other part serving as the control did not contain any meSIILCC. The 10-fold excess added was based the total protein concentration in the 37 °C aliquot of the DMSO-only set.

**Figure 7.**
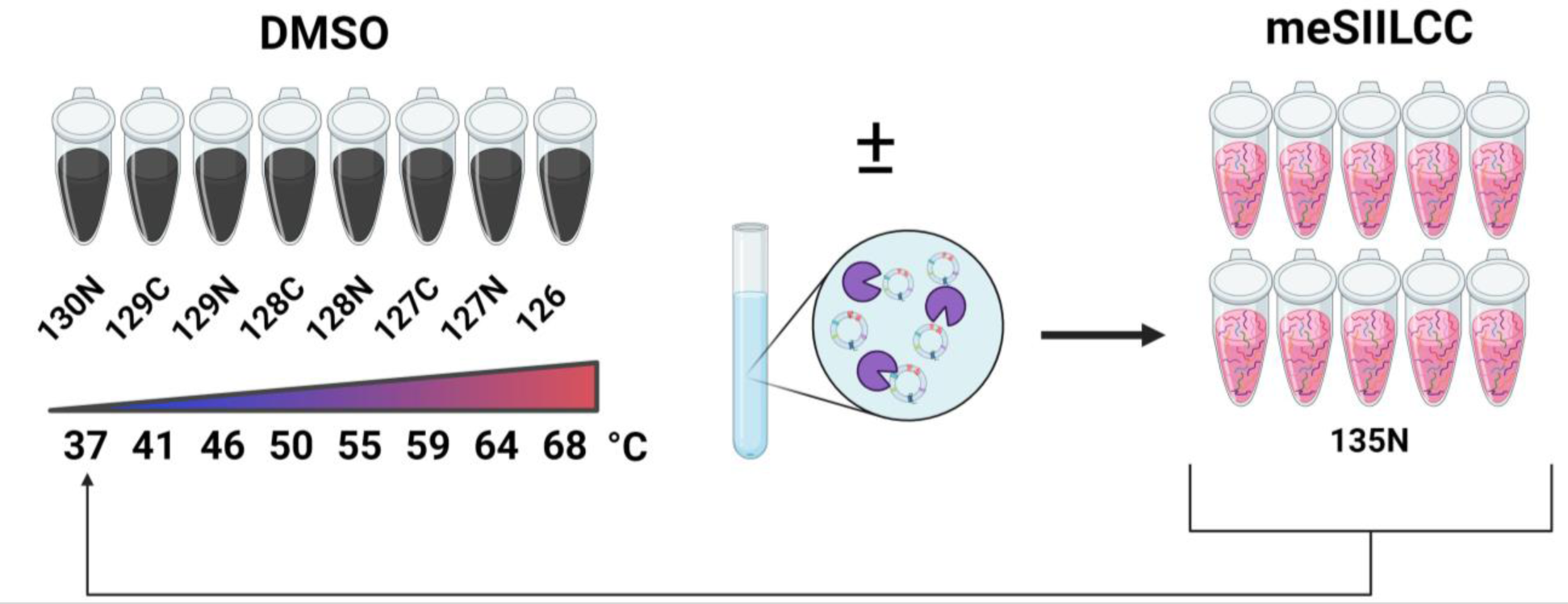
Isobaric labeling scheme for DMSO-only MS-TSA experiment. The first eight channels of TMTpro 18-plex was used for the heat-challenged intact cell aliquots. The meSIILCC (135N) was included at 10-fold the total protein content of the lowest heated aliquot. Created with www.Biorender.com

Figure 8 illustrates that multiple empty channels contain some level of isotopic interference, with the 134N channel containing the most significant interference (0.43 ±0.0484 level of normalized intensity). The contamination is the evident effect of 134N isotope impurities contained in the 135N reagent. Conversely, this level of 134N isotopic contamination was not detected in the 134N channel of the control sample that did include the meSIILCC component. To a lesser extent, the 130C, 131N, 133N, and 134C channels also show much lower levels of isotope contamination, which could be addressed by the implementation of correction factors.^13,36^

**Figure 8.**
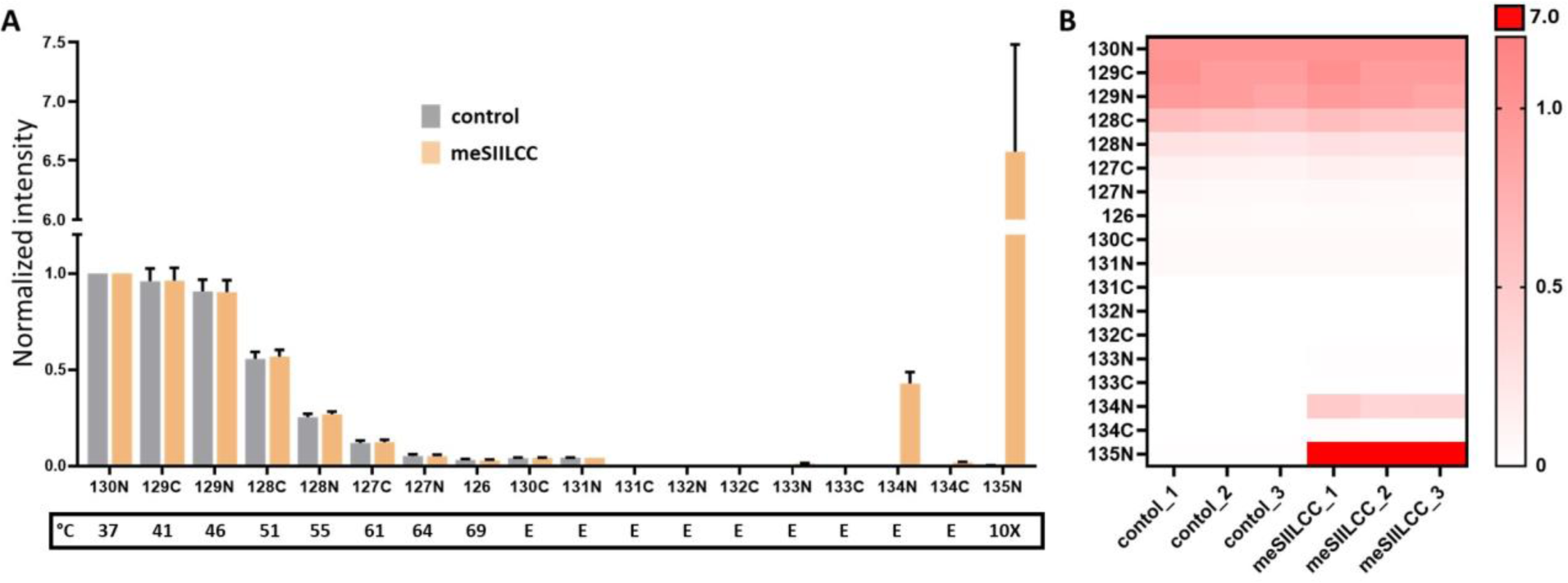
TMT reporter ion profiles for the DMSO only MS-TSA experiments for control and meSIILCC. **A)** Median reporter ion intensities for proteins detected across all replicates and found in both control and meSIILCC experiments. “E” represents those TMT channels left empty. **B)** Individual median reporter ion intensities for proteins detected across all replicates and found in both control and meSIILCC experiments.

### Over-Representation Analysis (ORA) indicates meSIILCC improves MP identifications in MS-TSA

To further assess the enhancement of MP identifications using meSIILCC, we performed ORA using the web-based functional enrichment analysis tool Enrichr^42,43^ using three subsequently defined data sets from the DMSO-only MS-TSA. A total of 5,436 protein groups that were identified across all replicates and found to be common between the control and meSIILCC experiments were used as the “common” dataset. We also identified 718 and 782 protein groups, which were also identified across all replicates, but were unique to the control and meSIILCC, respectively. The two unique sets were referred to as “unique control” and “unique meSIILCC” and were also subjected to ORA. We assumed that a baseline level of MPs was identified in conventional MS-TSA experiments,^20^ which would be captured by the common dataset, while additional MPs identified by the meSIILCC approach could be attributed to the meSIILCC-based enhancement of the MS-TSA approach. The limited number of MP-associated annotations from the unique proteins identified in the control set also underscores the advantages of a meSIILCC approaach; gene ontology (GO) terms containing “membrane” with p-values > 0.05 were counted, and 38, 17, and 3 terms in the common, unique meSIILCC, and unique control groups, respectively (Figure 9).

**Figure 9.**
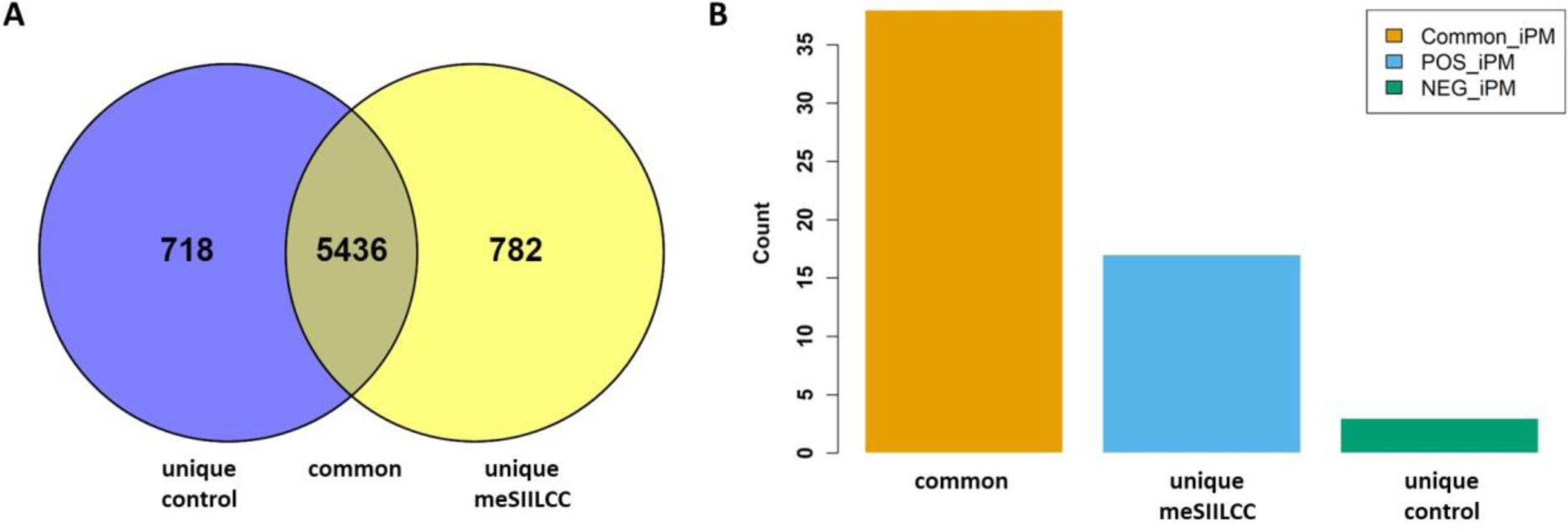
Protein identifications found across all replicates used to count cellular component GO terms containing “membrane”. **A)** The Venn diagram describing three datasets used for ORA, and **B)** Bar plot of cellular component GO terms, including the word “membrane” from the three defined datasets.

Next, we compared the top 10 GO terms based on p-value for the three datasets, which are shown in Table 1. In the common dataset, Intracellular Membrane-Bounded Organelle (GO:0043231), which includes organelles within a cell enclosed by a membrane, was found to be the most statistically significant (p-value: 1.26E-142). The GO term Nucleus (GO:0005634) was determined to have a p-value 1.54E-130, and Intracellular Non-Membrane-Bounded Organelle (GO:0043232), referring to organelles lacking a defining membrane was also shown to be highly statistically significant (p-value: 1.12E-63). Additionally, the Nuclear Lumen, Nucleolus, and Mitochondrial Matrix were significantly represented in the common group.

**Table 1.**
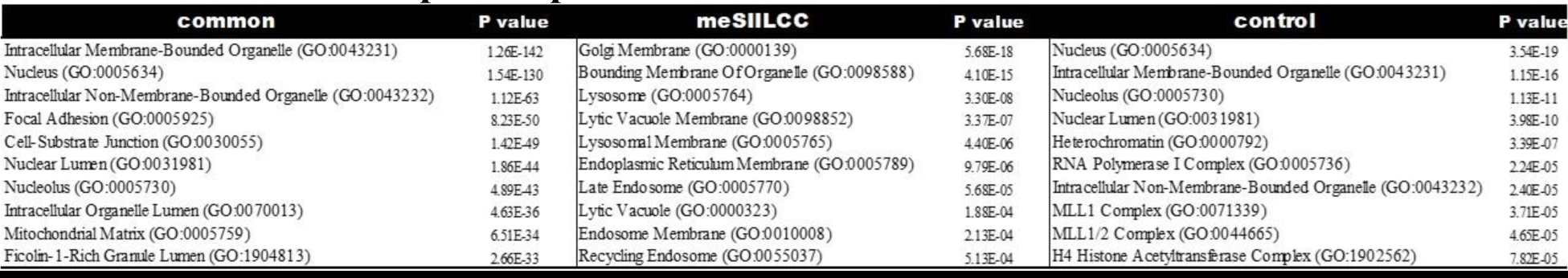
Top 10 cellular component GO terms for common, unique meSIILCC, and unique control datasets with respective p-values.

Cellular component GO terms associated with nuclear structures or intracellular organelles (including proteins contained within these organelles) were identified in the unique control dataset; Nucleolus (GO:0005730, p-value 1.13E-11) and Heterochromatin (GO:0000792, p-value 3.39E-07).

The unique meSIILCC group displayed a marked over-representation in membrane-associated terms: a highly statistically significant p-value (5.68E-18) was determined for the GO term Golgi Membrane (GO:0000139), which would include proteins found within that membrane. Many other highly statistically significant GO terms associated with membranes were also identified: Bounding Membrane of Organelle (GO:0098588), including membranes encasing cellular organelles (p-value 4.10E-15); Lysosome (GO:0005764) and Lysosomal Membrane (GO:0005765) having p-values of 3.30E-08 and 4.40E-06, respectively. The Endoplasmic Reticulum Membrane (GO:0005789) and Endosome Membrane (GO:0010008) that relates to proteins annotated to these membranes were also found to have significant p-values. The high abundance of cellular component GO terms for proteins identified in the unique meSIILCC dataset was also indicative of MP-enriched dataset.

### An increase in the number of peptides per protein for proteins annotated with the term “plasma membrane” that were identified in meSIILCC MS-TSA experiments

We hypothesized that using the meSIILCC approach would increase not only the number of MP identifications, but also the number of identified peptides per protein, with the effect of improved data quality^44–46^. To minimize potential bias coming from the highly stochastic precursor sampling in data-dependent acquisition (DDA)^47,48^, we excluded identified proteins unique to the meSIILCC experiment. The analysis was limited to proteins identified in the common set, which corresponded to 48,540 (±231) and 46,825 (±2,032) unique peptides for the control and meSIILCC conditions, respectively. The previously described UniProt-IPA database was then used to filter the peptide lists by selecting for the “plasma membrane” (PM) subcellular localization, which resulted in a subset of 496 proteins from the common dataset. The PM-annotated subset corresponded to 3,807 (±22) and 4,866 (±241) unique identified peptides for the control and meSIILCC groups, respectively.

Unique peptide identifications per protein from the common and PM-annotated datasets for meSIILCC and control were overlaid as violin plots, as shown in Figure 10. The plots for the common dataset closely overlapped, with median peptides per protein of 6.3 and 6.0 for control and meSIILCC, respectively, while plots for the PM-annotated subset showed a marked increase in identified peptides per protein using the meSIILCC approach, with medians of 4.7 and 6.3 for control and meSIILCC, respectively, demonstrating the positive effect of the meSIILCC approach.

**Figure 10.**
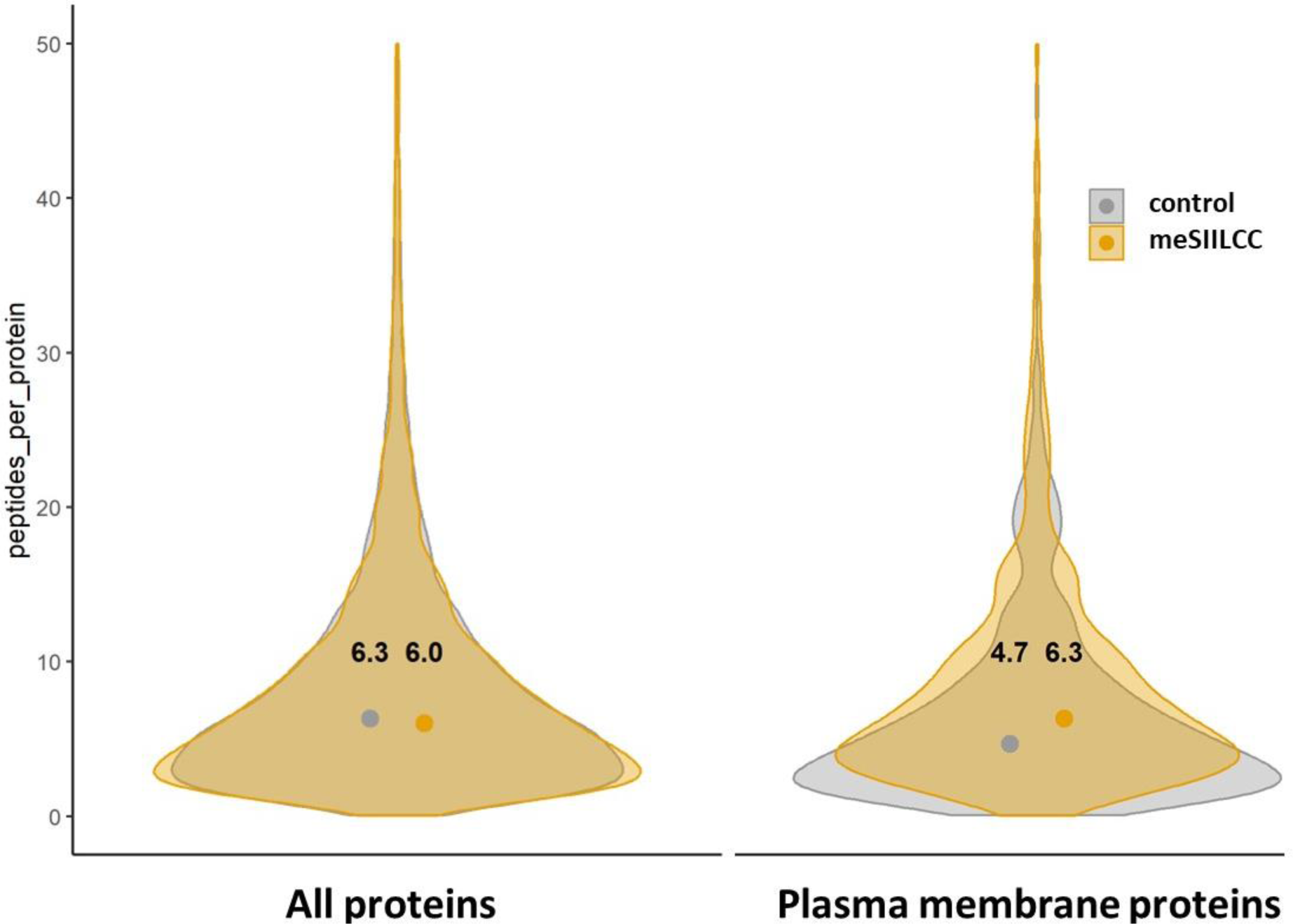
Violin plot of peptides per protein from the common data set for **A)** All and **B)** PM-annotated proteins.

### Protein T_m_ from DMSO curves were found to be similar in the meSIILCC and control MS-TSA experiments

To further assess the impact of meSIILCC on T_m_, we first compared the linear relationship of T_m_ from control and meSIILCC MS-TSA experiments. Reporter ion intensities for proteins from the previously described common dataset and PM-annotated subset were used to determine T_m_. The T_m_ values were determined at a half-maximal normalized intensity and visualized by a scatter plot, and those outside the measured range (37-68.5 °C) were removed, resulting in 4,512 and 409 paired T_m_ values from the common dataset and PM-annotated subsets, respectively.

In the common dataset, median T_m_ values were determined to be 51.98 °C for the control and 51.95 °C for meSIILCC, while in the PM annotated subset, median T_m_ values were 53.69 and 53.46 °C for the control and meSIILCC datasets, respectively (Figure 11 A-B). Notably, these findings align with prior work^20^, where an increase in T_m_ for MP for intact MS-TSA experiments was observed.

**Figure 11.**
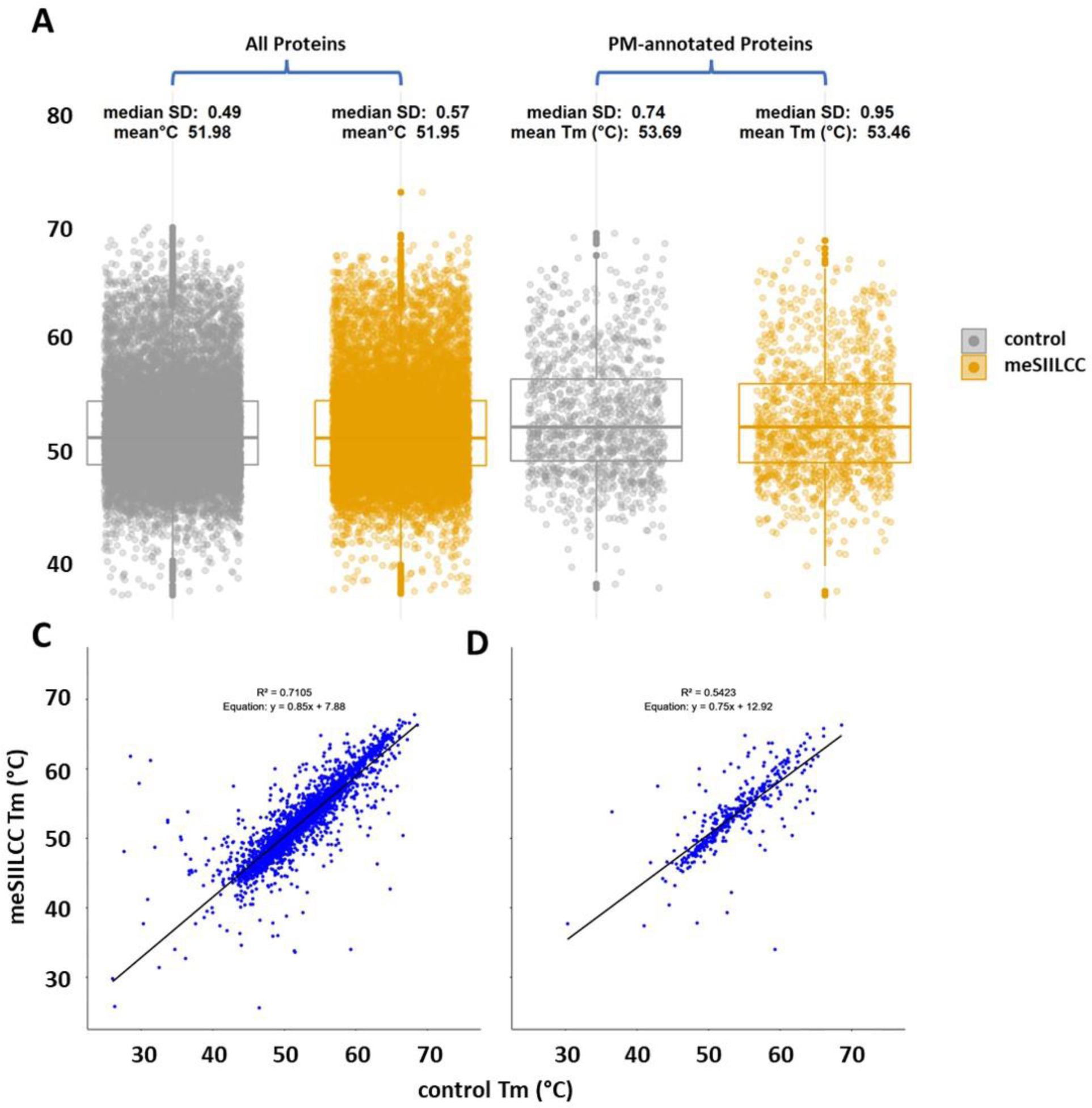
Comparison of T_m_ determined in the common and PM-annotated datasets for control and meSIILCC. **A)** Scatter plot of T_m_ for the common dataset and **B)** PM-annotated subset. **C)** Linear fitting of T_m_ in the common dataset and **D)** PM-annotated subset.

The coefficient of determination (R^^2^) was calculated as 0.7105 and 0.5423 for the control dataset and PM annotated subset (Figure 11 C-D), respectively, indicating strong linear relationships in both the common dataset and PM-annotated subset.

### Conclusion

In this work, we successfully developed a membrane-enriched stable isotope isobarically labeled carrier channel approach for mass spectrometry-based thermal stability assay (MS-TSA) to improve the detection and quantification of membrane proteins in MS-TSA experiments.

An initial POC study demonstrated improvements across all proteomic identifications with the exception of MS^2^ scans, with a 10% increase in identifications of proteins annotated as MPs. Using meSIILCC, 2.6-fold more peptides were exclusively identified compared to the control, and a hydrophobicity index demonstrated a decrease in median GRAVY values; −0.671 for the peptides exclusively found in the control, compared to −0.371 for those exclusively found in the meSIILCC group. The increased GRAVY score suggests that a more hydrophobic set of peptides was identified in the meSIILCC group, which is characteristic of MPs.

To further improve the meSIILCC approach, four MP-enrichment techniques were assessed. We identified a superiorly performing MP-enriching protocol using the iPM extract from Invent Biotechnology’s Minute™ Kit. The highest number of positive log_2_FC values annotated with PM (61%), in combination with the lowest amount of log_2_FC values annotated with subcellular locations other than PM were identified in the iPM extract.

In a DMSO-only MS-TSA experiment, the optimally performing meSIILCC was labeled with the TMT 135N reagent and spiked in at a 10-fold protein amount ratio based on the total protein amount in the lowest-heated aliquot. The 130C-134C TMT channels were left empty to evaluate isotope impurity interference, and a significant amount of interference (0.43 ±0.0484 normalized intensity) was observed in the 134N channel and to a lesser extent in the 130C, 131N, 133N, and 134C channels.

An ORA was conducted on three datasets defined from the DMSO-only MS-TSA experiment, and 17 cellular component GO terms containing “membrane” in the term definition with p-values > 0.05 were found in the unique meSIILCC dataset, compared to 3 in the unique control dataset. Additionally, we found that all of the top ten cellular component GO terms for proteins identified in the unique meSIILCC dataset specifically describe the membrane origin of the protein or, in the context of describing an entire organelle, imply the presence of a membrane.

We found that a common set of PM-annotated proteins between the control and meSIILCC groups for the DMSO-only MS-TSA, which were identified across all replicates, demonstrated an increase in peptides per protein when using meSIILCC. The median values of peptides per protein for PM-annotated proteins in the DMSO-only MS-TSA experiment were 4.7 and 6.3, respectively.

Protein T_m_ in the DMSO-only MS-TSA were found to be similar and having a linear relationship in the meSIILCC and control MS-TSA experiments. The presence of meSIILCC did not significantly affect T_m_ values when using 126-130N TMTpro reagents, and further experiments are ongoing to demonstrate that when using the 135N channel for meSIILCC, all channels except the 134N can be used in the TMTpro 18-plex for MS-TSA experiments. We also are planning to benchmark our meSIILCC-aided approach compared to cs-TPP.

